# M-DeepAssembly2: A Web Server for Predicting Multiple Conformations of Multi-domain Proteins Using Deep Learning

**DOI:** 10.1101/2025.10.23.684122

**Authors:** Xinyue Cui, Lingyu Ge, Xinguang Yang, Xuhui Li, Guijun Zhang

## Abstract

The biological functions of proteins often depend on dynamic transitions between multiple conformational states, and for multi-domain proteins, inter-domain dynamics play a critical role in their biological functions. Despite significant advances in static structure prediction, accurately modeling these multiple conformations remains a major challenge. Here, we present M-DeepAssembly2, a web server designed for modeling multiple conformations of multi-domain proteins using deep learning-predicted inter-domain and intra-domain constraints. The server operates in three stages. Firstly, evolutionary, physicochemical, and geometric features are extracted from the input and fed into AlphaFlex, a flexible residue prediction network. This process generates multiple distance maps that effectively capture both inter-domain and intra-domain interactions. Secondly, supplementary distance maps that characterize inter-domain interactions are generated using the DeepAssembly method. Finally, all distance maps are integrated as constraints in a multi-objective protein optimization to generate multiple conformational states of the protein. Compared to its predecessor, M-DeepAssembly2 leverages a flexible residue prediction network that directionally decouples coevolutionary information to generate heterogeneous distance maps. Benchmark results demonstrate that this strategy enables accurate prediction of protein multiple conformations and significantly improves the precision of static multi-domain protein structure prediction.

## Introduction

In nature, most proteins comprise two or more structural domains. These domains serve as fundamental evolutionary and functional units that exhibit considerable conformational flexibility. Changes in their relative orientation often dictate the dynamic behaviors and regulatory mechanisms of molecular machines, signaling proteins, and allosteric proteins^1^. Recent advances in deep learning have markedly accelerated progress in protein structure prediction. End-to-end methods such as AlphaFold^2,3^, RoseTTAFold^4^, and ESMFold^5^ have largely addressed the long-standing challenge of achieving high-accuracy prediction of static monomeric structures^6^. However, these methods are primarily trained on and optimized for single, static conformations and thus remain limited in capturing the dynamic inter-domain rearrangements and interactions of multi-domain proteins under physiological conditions. Consequently, there is an urgent need for new methods to characterize the multiple conformational states of multi-domain proteins, which will provide important mechanistic insights into their functional dynamics and allosteric regulation.

The breakthrough in static structure prediction has spurred the development of methods for characterizing the dynamic conformational landscapes of proteins, which can be broadly categorized into experimental and computational methods. Experimental methods such as nuclear magnetic resonance (NMR)^7^, cryo-electron microscopy (cryo-EM)^8^, and X-ray crystallography^9^ provide high-resolution structural information, but their applications are limited by stringent crystallization requirements and the inherent challenges posed by sparse, ambiguous, or noisy data^10^. Computational methods include molecular dynamics (MD) simulation, Monte Carlo (MC) sampling, and deep learning-based methods. MD simulations offer high temporal resolution of protein dynamic changes by accurately characterizing atomic interactions, but they are computationally expensive and strongly dependent on the accuracy of force fields^11^. Deep learning-based methods can be further divided into AlphaFold-based extensions and generative models. Specifically, representative methods are introduced as follows: AF-Cluster^12^ predicts multiple conformations by clustering multiple sequence alignments (MSAs), while AFsample2^13^ employs column-masking strategies within MSAs to explore the dynamic properties of proteins. Generative models such as AlphaFLOW^14^ leverage flow-matching algorithms to construct conformational ensembles, whereas methods like DiG^15^ and BioEmu^16^ exploit diverse datasets to enable large-scale investigations of equilibrium distributions and conformational transitions. It should be noted that these methods are inherently data-driven, and their performance depends critically on the conformational coverage of the training data. In the context of scarce dynamic data, the strategy based on MC sampling demonstrates obvious advantages. Multi-population protein conformation sampling algorithms can sample a wide range of conformational states under the guidance of multiple energy functions^17^. Similarly, iterative exploration strategies integrating multi-objective optimization, geometric refinement, and structural similarity clustering have shown considerable promise in conformational sampling^18^. Despite these advancements, the precise modeling of protein multiple conformations poses a significant challenge, particularly for multi-domain proteins, with considerable room for improvement in this field.

In our previous work on multi-domain protein modeling, M-DeepAssembly^19^, we extracted intra-domain and inter-domain information from proteins and incorporated them as constraints in a multi-objective protein conformation sampling algorithm, enabling comprehensive exploration of conformational space and high-accuracy modeling of multi-domain proteins. Although this method partially alleviates the challenges caused by weakened coevolutionary information, it still fails to capture the multiple conformational states of proteins.

This limitation arises because the spatial constraints are derived from static structure predictions and therefore cannot adequately reflect the diversity of protein conformations. Consequently, building upon M-DeepAssembly, it is essential to introduce multiple constraints that incorporate diverse inter-domain and intra-domain information for achieving accurate modeling of protein multiple conformations.

In this work, we proposed M-DeepAssembly2, an upgraded version designed to model multiple conformations of multi-domain proteins. Compared to its predecessor, M-DeepAssembly2 utilizes the flexible residue prediction network AlphaFlex^20^ to adjust inter-domain and intra-domain interactions within the distance maps selectively. These modified distance maps are then integrated alongside the DeepAssembly-predicted inter-domain map to serve as constraints in a multi-objective optimization model, thereby enabling the modeling of multiple conformational states for multi-domain proteins. The benchmark results demonstrate that M-DeepAssembly2 achieves superior performance in multi-conformation prediction tasks and substantially outperforms its predecessor in static multi-domain structure prediction. These results indicate that M-DeepAssembly2 can serve as a valuable tool for investigating the complex three-dimensional structures and dynamic mechanisms of multi-domain proteins.

## Results

### Overview of the M-DeepAssembly2

M-DeepAssembly2 is a web server designed for modeling multiple conformations of multi-domain proteins. As shown in **Figure 1**, users can submit either a full-length sequence, individual domain sequences, or a combination of individual domain sequences and structures. For full-length sequence submissions, M-DeepAssembly2 utilizes DomBpred^21^ to split them into individual domain sequences. Any individual domain sequences without provided structures are processed by AlphaFold3^3^ to generate the corresponding structures. Next, evolutionary features (MSA embeddings^22^, BLOSUM62 substitution matrix^23^), physicochemical features (Rosetta energy terms^24^, solvent accessibility), and geometric features (ultrafast shape recognition (USR)^25^, orientation) are extracted. These features are input into AlphaFlex^20^, a flexible residue prediction network, which generates distance maps richly encoding both inter-domain and intra-domain interactions. In parallel, we employed our in-house DeepAssembly^26^ model to predict an additional, singular distance map specifically characterizing inter-domain interactions. By integrating these complementary distance maps, we developed a multi-objective optimization model and designed a Metropolis Monte Carlo (MMC) sampling scheme guided by Pareto optimization. Through performing multiple iterative refinement cycles, the framework ultimately achieves efficient exploration and generation of diverse conformational states for multi-domain proteins.

**Figure 1.**
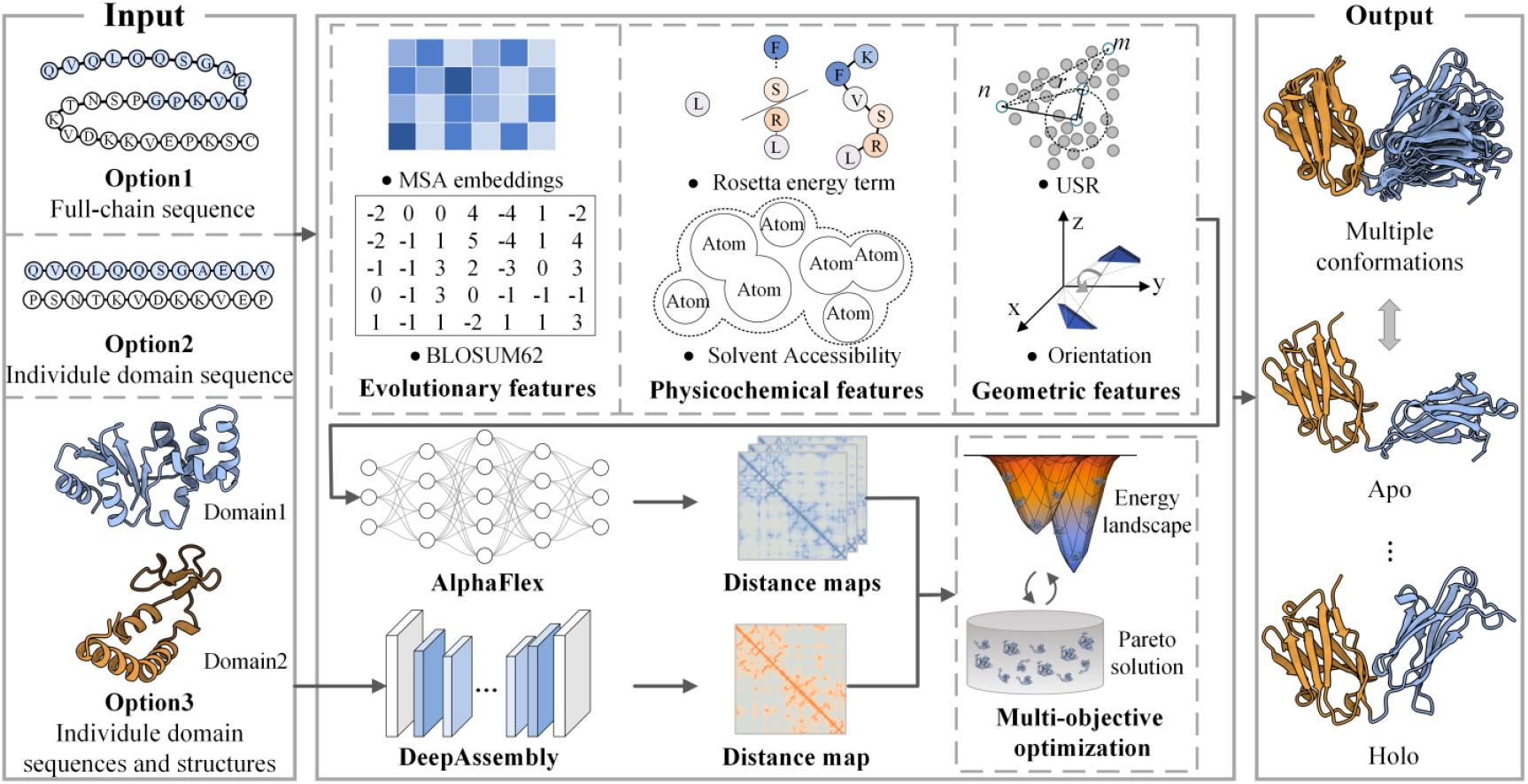
Overview of M-DeepAssembly2. The method takes as input protein sequences or structures and outputs multiple conformational states. Users can provide one of three input options: a full-length sequence, individual domain sequences, or a combination of individual domain sequences and structures. Full-length sequences are first segmented into domains using DomBpred^21^, and domain sequences without structures are processed by AlphaFold3^3^ to predict their corresponding structures. Subsequently, evolutionary, physicochemical, and geometric features extracted from the input are fed into the AlphaFlex^20^ to generate both inter-domain and intra-domain distance maps, while an additional inter-domain distance map is generated using DeepAssembly^26^. These distance maps are subsequently integrated into a multi-objective optimization framework to predict multiple conformations.

### Performance on multiple conformations prediction

To evaluate the performance of M-DeepAssembly2 in modeling multiple conformations of multi-domain proteins, we constructed a test set based on 91 commonly used Apo/Holo pairs from the CoDNaS^27^ dataset. Among these, 14 pairs were identified as multi-domain targets, which were ultimately used as the benchmark for evaluation (see **Methods** for more details). The similarity between the predicted conformations and the native Apo/Holo states was systematically evaluated using template modeling score (TM-score)^28^ and root mean square deviation (RMSD). We compared our method against two representative state-of-the-art baseline approaches: AF-Cluster^12^, which employs MSA clustering techniques based on AlphaFold^2^, and AlphaFLOW^14^, which utilizes flow-matching generative models. The benchmarking results demonstrate that M-DeepAssembly2 outperformed the reference methods in 85.7% of targets for the Apo state. For the Holo state, it surpassed AF-Cluster and AlphaFLOW in 64.3% and 71.4% of targets, respectively (**Figure 2a-d, Supplementary Table S1 and S2**). This advantage is confirmed by the average TM-scores (**Figure 2e**). Our predicted Apo and Holo states achieved average TM-scores of 0.91 and 0.86, respectively. These scores correspond to a significant improvement of 13.75% (Apo) and 6.2% (Holo) over AF-Cluster, as well as an enhancement of 4.6% (Apo) and 3.6% (Holo) over AlphaFLOW. Furthermore, this advantage was further substantiated by superior RMSD values (**Figure 2f**).

**Figure 2.**
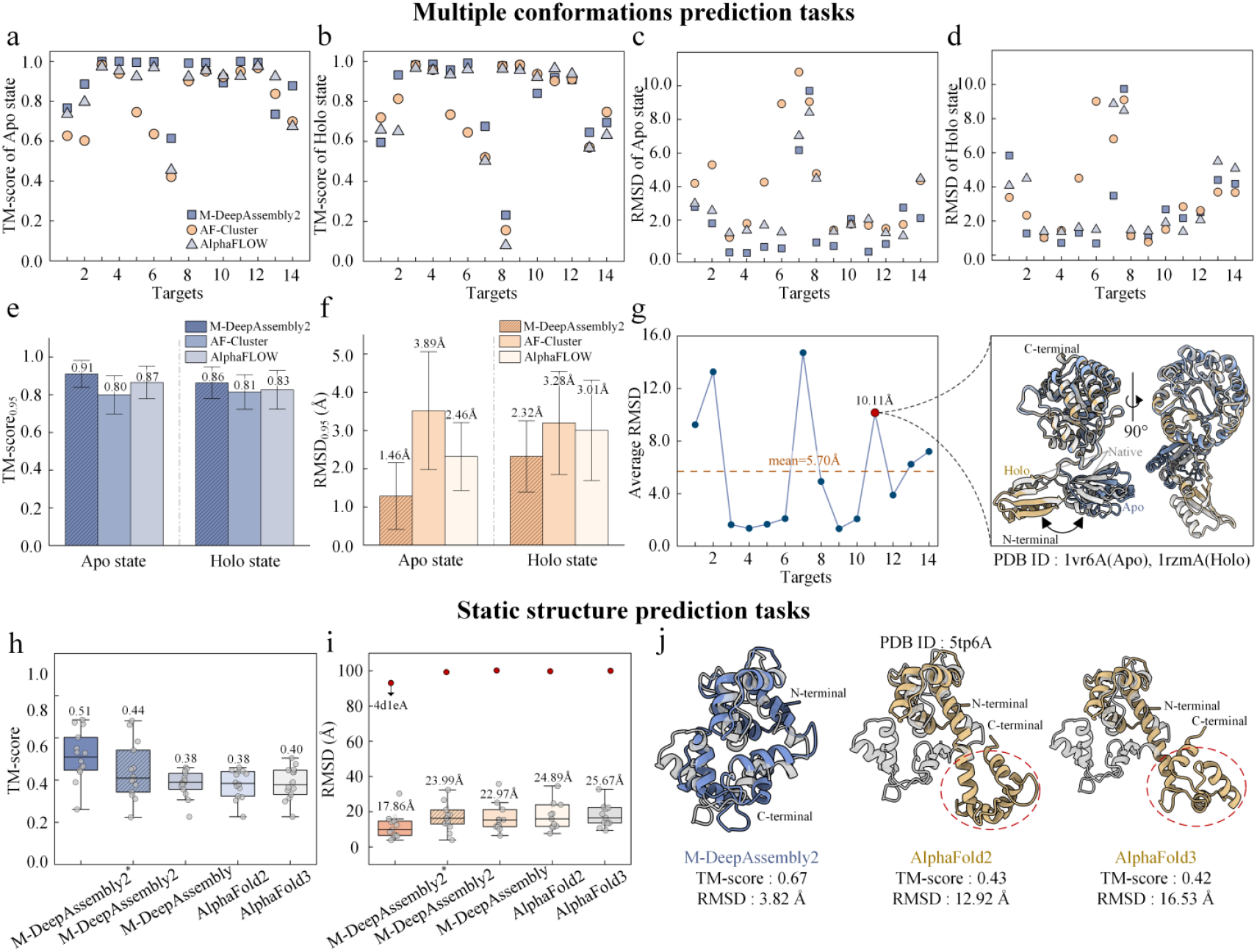
Performance evaluation of M-DeepAssembly2 in multiple conformations and static structure prediction tasks. **a-f**, Performance comparison of M-DeepAssembly2 with AF-Cluster^12^ and AlphaFLOW^14^ in multi-conformation prediction. a-d show the comparison based on TM-score and RMSD metrics, while e-f present the average performance of different methods. The bar height indicates the mean; error bars indicate 95% confidence intervals. **g**, RMSD distribution between the native Apo and Holo states in the multi-conformation test set. The right panel shows a representative example of Apo and Holo state predictions for 1vr6A by M-DeepAssembly2. The native crystal structure is shown in gray, while M-DeepAssembly2’s predicted Apo state is shown in yellow and its predicted Holo state is shown in blue. **h-i**, Performance evaluation in static structure prediction, which shows the average TM-score (h) and RMSD (i) comparisons among M-DeepAssembly2, M-DeepAssembly^19^, AlphaFold2^2^, and AlphaFold3^3^. M-DeepAssembly2* refers to the predicted structure exhibiting the highest structural similarity to the native structure. M-DeepAssembly2 denotes the best model selected using our in-house developed MViewEMA^29,30^ model quality assessment algorithm. **j**, The example of the calmodulin variant 34 (CaM34, PDB ID: 5tp6A) predicted by different methods. The native structure is colored gray. The blue structure represents the prediction from M-DeepAssembly2, while the yellow structures are the predictions from AlphaFold2 and AlphaFold3. All M-DeepAssembly2 results were obtained by assembling individual domain structures predicted using AlphaFold3.

As a representative case, we examined the Phospho-2-dehydro-3-deoxyheptonate aldolase (DAHP synthase) from Thermotoga maritima. The Apo structure (PDB ID: 1vr6A) captures the unliganded state of the enzyme, whereas the Holo structure^31^ (PDB ID: 1rzmA) represents a complex bound to metal cofactors and substrate analogues. The two conformational states exhibit a significant rotation of the N-terminal domain (**Figure 3g**), with the difference quantified by a TM-score of 0.81 and an RMSD of 10.11 Å (**Supplementary Table S3**). Notably, M-DeepAssembly2 successfully captured these two conformations, achieving predicted TM-scores of 0.99 for the Apo state and 0.92 for the Holo state. These outstanding performances can be primarily attributed to the incorporation of the flexible residue prediction network (AlphaFlex^20^). This network enables the characterization of inter-domain and intra-domain interaction features in multi-domain proteins at the structural level, thereby generating distance maps that capture conformational heterogeneity. This approach significantly expands the conformational sampling space, allowing the model to explore a broader range of structurally plausible states with higher accuracy.

**Figure 3.**
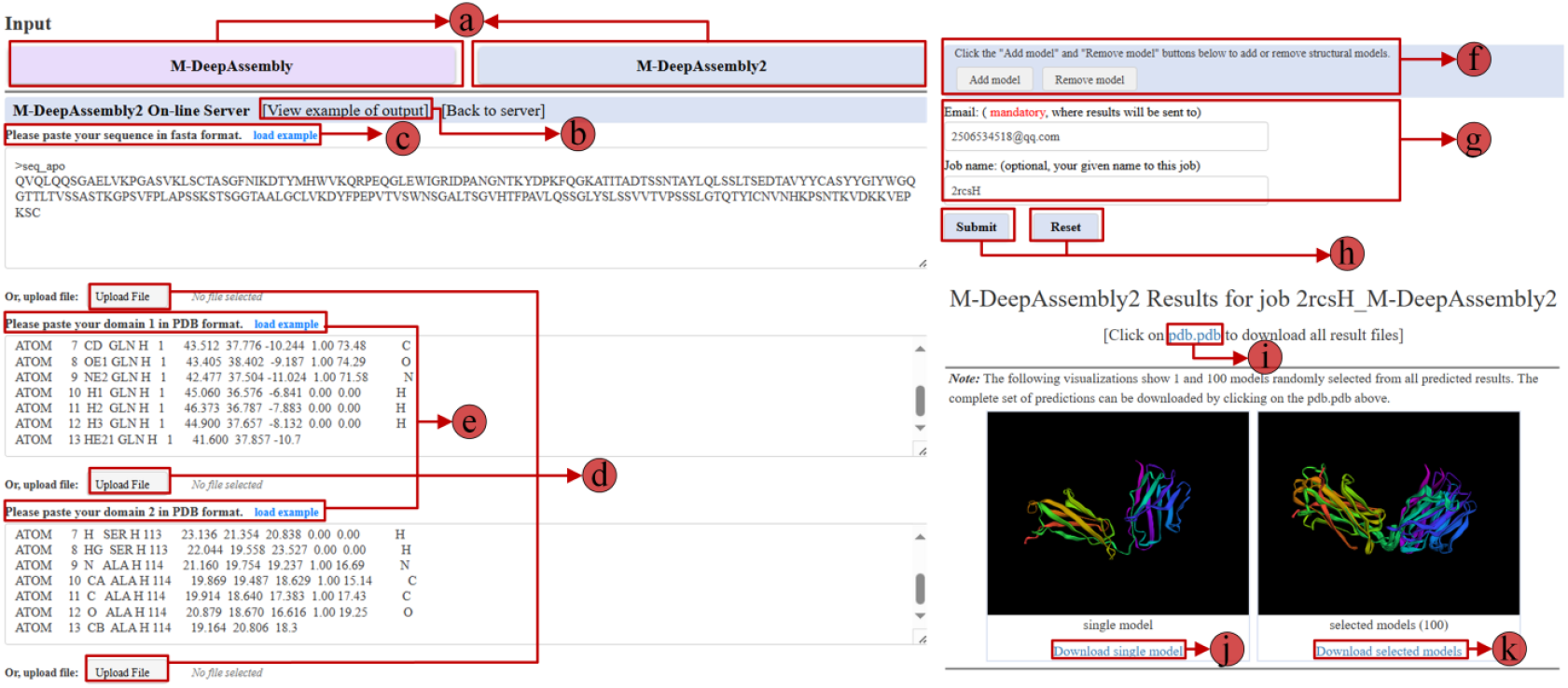
Input and results interfaces of the M-DeepAssembly2 web server. **a**, Options for multi-domain protein static structure prediction (M-DeepAssembly^19^) and multiple conformations prediction (M-DeepAssembly2); **b**, Example display; **c**, Target sequence loading box; **d**, local file loading button; **e**, Target individual domain structure loading box; **f**, Add or remove domain loading box **g**, Email address and job name box; **h**, Submit and reset buttons; **i**, Download link for the complete set of predicted structures. **j**, Visualization of a single, randomly selected predicted structure. **k**, Visualization of an ensemble comprising 100 randomly selected predicted structures.

### Performance on protein static structure prediction

To assess the capability of M-DeepAssembly2 in multi-domain protein static structure prediction, we evaluated its performance on a benchmark dataset comprising 14 multi-domain proteins selected from the original M-DeepAssembly^19^ test set (see **Methods** for details). These targets represent particularly challenging cases for previous methods, as indicated by their best predicted TM-scores being below 0.5 (see **Supplementary Table S4** for details). The set includes proteins with 2-4 contiguous domains (88-894 amino acids). For each protein, individual domains were first predicted by AlphaFold3^3^ and then assembled using M-DeepAssembly2. We compared the assembled structures with those obtained from the previous version of M-DeepAssembly^19^, AlphaFold2^2^, and AlphaFold3^3^, using TM-score^28^ and RMSD as evaluation metrics for the prediction results.

To ensure clarity in performance evaluation, we distinguished two types of predicted results. M-DeepAssembly2* denotes the prediction with the highest structural similarity to the native structure among outputs, representing the upper bound of the method’s practical predictive performance. M-DeepAssembly2 refers to the best predicted result selected by our in-house model quality assessment algorithm (MViewEMA^29,30^), reflecting the method’s practical predictive performance. As illustrated in **Figure 2h-i**, the test results show that M-DeepAssembly2 demonstrated superior performance in multi-domain protein static structure prediction. The average TM-score of M-DeepAssembly2 reached 0.44 (**Figure 2h, Supplementary Table S4**), representing improvements of 15.8%, 15.8%, and 10% over M-DeepAssembly^19^, AlphaFold2^2^, and AlphaFold3^3^, respectively. Furthermore, M-DeepAssembly2 also achieved a lower average RMSD of 23.99 Å compared to AlphaFold2 and AlphaFold3 (**Figure 2i**). Notably, M-DeepAssembly2* achieved the highest performance (**Figure 2h-i, Supplementary Table S5**), surpassing both M-DeepAssembly2 and all other benchmark methods. These results demonstrate that there remains room for improvement in the model quality assessment used for selecting the best predictive structure.

It’s worth noting that all methods performed poorly on the 4d1eA target. Structural analysis revealed that this protein consists of multiple linearly arranged contiguous domains (**Supplementary Figure S1**). This spatial arrangement may significantly attenuate inter-domain interactions, increase conformational flexibility, and thereby challenge predictive models in accurately capturing the relative orientations of the domains. Nevertheless, M-DeepAssembly2 still demonstrated superior performance on the test set. A specific case that highlights the effectiveness of our method is calmodulin variant 34 (CaM34) in complex with the iNOS CaM binding domain peptide^32^ (**Figure 2j**, PDB ID: 5tp6A). While end-to-end methods like AlphaFold2 and AlphaFold3 exhibited significant deviations in the C-terminal domain’s orientation, M-DeepAssembly2 successfully captured the correct inter-domain orientation by assembling domains predicted by AlphaFold3, resulting in an improved TM-score of 0.67. These results, particularly the significant improvements over the previous version of M-DeepAssembly^19^, further validate the efficacy of the newly introduced flexible residue prediction network. This network significantly enhances the accessible conformational space of proteins. When integrated with a multi-objective optimization algorithm, it facilitates higher-precision modeling of static structures for multi-domain proteins.

### Webserver interface

#### Server input

The M-DeepAssembly2 web server offers both multiple conformations and multi-domain protein static structure prediction modeling services, allowing users to choose according to their needs. As illustrated in **Figure 3**, the server accepts protein sequences (full-chain or individual domain sequences) in FASTA format or individual domain structures in PDB format. Users can either paste the corresponding information or upload local files directly. For assemblies involving more than two domains, additional domains can be added using the “Add model” button, while incorrectly added domains can be removed via the “Remove model” button. If the user only has a protein sequence, it’s recommended to utilize state-of-the-art protein structure prediction tools to obtain individual domain structures. However, this step is not mandatory. When only a full-chain sequence is submitted, the server automatically performs domain parsing using our in-house DomBpred^21^ method. Subsequently, all individual domain sequences are processed by AlphaFold3^3^ to predict their corresponding three-dimensional structures. Users must provide a valid email address to receive completion notifications. In addition, multiple submissions can be managed and tracked using custom job names. After completing all required inputs, clicking the “Submit” button triggers an email address confirmation window (see **Supplementary Figure S2**). Once the email address is confirmed, the job is successfully submitted, and the server automatically sends a job submission confirmation email to the user.

#### Server output

The M-DeepAssembly2 web server provides a user-friendly and efficient workflow. Upon job submission, the webpage automatically redirects to the results page, which refreshes every five seconds to provide real-time updates on task progress (see **Supplementary Figure S3**). Once the job is complete, the server sends a notification email containing a download link, which includes a submission timestamp for convenient tracking and distinction between multiple jobs. The results page (see **Figure 3**) displays the job name along with a visualization of a randomly selected predicted structure for rapid inspection. Users are provided with flexible download options: the complete ensemble of predicted structures can be retrieved collectively via the “pdb.pdb” link (**Figure 3i**), or selected models can be downloaded using the “Download selected models” (**Figure 3k**) functionality. It is important to note that the server is optimized for multi-domain proteins with sequence lengths less than 1024 amino acids. While prediction for medium-sized proteins (300-500 residues) is typically completed within 12 hours, the runtime may be extended under heavy job load.

## Discussion

In this study, we introduce M-DeepAssembly2, a web server that significantly advances the computational modeling of multi-domain proteins by integrating a flexible residue prediction network with a multi-objective protein conformation sampling algorithm. This strategy can accurately predict both multiple conformational states and static structures of multi-domain proteins. We rigorously evaluated the efficacy of our method on a benchmark dataset comprising 14 Apo/Holo protein pairs and an additional 14 static multi-domain proteins. The results clearly demonstrate M-DeepAssembly2’s superiority over benchmark methods. Specifically, our model outperformed the comparison method on 12/14 of the tasks for Apo state prediction. Notably, for the more challenging Holo state prediction, which involves ligand-induced conformational changes. Even in the absence of explicit ligand information, M-DeepAssembly2 still outperformed AF-Cluster^12^ and AlphaFLOW^14^ on 9/14 and 10/14 tasks, respectively. This performance highlights our model’s unique ability to effectively capture and model the intricate conformational changes that occur upon ligand binding. Furthermore, the assembly of multi-domain proteins by M-DeepAssembly2 showed a remarkable 15.8% improvement in accuracy over the original M-DeepAssembly, further confirming the enhanced performance of its static structure prediction.

Despite these significant advancements, there remains considerable room for further improvement. Firstly, our current model, which assembles domains as rigid units, effectively captures large-scale inter-domain motions but is inherently limited in its ability to simulate subtle conformational fluctuations within individual domains. Given that these fine-grained dynamics are critical for regulating protein function, future work should focus on developing advanced methods capable of simultaneously modeling both large-scale domain rearrangements and fine-grained inter-domain and intra-domain structural fluctuations. Secondly, the interaction modes between domains are highly analogous to the mechanisms of inter-chain interactions. Building on this insight, the core algorithms of M-DeepAssembly2 can be further generalized to model the multiple conformational states of protein complexes. Continued efforts along these directions will enable M-DeepAssembly2 as a robust server for revealing the dynamic nature of protein function.

## Materials and Methods

### Dataset

To evaluate the performance of our model on the protein multiple conformations prediction task, we selected 91 Apo/Holo protein pairs from the CoDNaS^27^ database. We first used our in-house developed DomBpred^21^ tool to identify 37 multi-domain pairs. After excluding 23 pairs with discontinuous domains, a final set of 14 Apo/Holo pairs was retained as the test set (including 12 two-domain and 2 three-domain proteins; for details, see **Supplementary Table S1, S2, and S3**). The Apo and Holo states in this test set exhibit significant conformational differences, with an average RMSD of 5.70 Å (**Supplementary Figure S4**). Additionally, to further assess our model’s performance on static structure prediction of multi-domain proteins, we selected 14 challenging targets with TM-scores < 0.5 from the M-DeepAssembly^19^ benchmark dataset as a supplementary test set (for details, **see Supplementary Table S4**).

### Benchmark methods

For the multi-conformational structure prediction task, AF-Cluster^12^ and AlphaFLOW^14^ were both executed using the default parameters provided in their official GitHub repositories (https://github.com/HWaymentSteele/AF_Cluster; https://github.com/bjing2016/alphaflow). The number of conformations generated by AF-Cluster corresponds to the number of MSA clusters produced by the method itself, while AlphaFLOW generated 1,000 conformations. For consistency, our method was also used to generate 1,000 conformations for comparison. For the multi-domain protein static structure prediction task, the predictions of M-DeepAssembly^19^ were obtained directly from its original publication. The results of AlphaFold2^2^ were produced using the publicly available GitHub implementation with default parameters (https://github.com/google-deepmind/alphafold), and the results of AlphaFold3^3^ were obtained through its official web server (https://alphafoldserver.com/?golgi=true).

### Input features

The M-DeepAssembly2 integrates evolutionary, physicochemical, and geometric features to comprehensively characterize the input protein sequence and structure, thereby enabling the development of a multiple conformation prediction algorithm. For evolutionary features, we obtained embeddings from MSAs using the protein language model ESM-MSA-1b^22^, which was generated from the UniRef90^33^, UniRef30^34^, and BFD^35^ databases (represented as L×L×144 and L×768 tensors). We also incorporated the BLOSUM62 substitution matrix^23^ (L×24) to quantify amino acid similarities. The physicochemical features include both one-body (p_aa_pp, rama_prepro, omega, fa_dun; L×L×7) and two-body (fa_atr, fa_rep, fa_sol, lk_ball_wtd, fa_elec, hbond_bb_sc, hbond_sc; L×7) Rosetta energy terms^24^. The solvent accessible surface area (L×1) was also included as an additional feature. For geometric features, we used USR (L×3) and orientation (L×L×6) to quantify the local and global topology of the protein.

### Flexible residue prediction network

The Flexible residue prediction network, based on a Transformer architecture, learns the intrinsic flexibility of amino acids from both static PDB and dynamic MD simulation databases. This enables it to identify flexible regions within a given structure accurately. By utilizing the predicted flexibility residue index (The top 5% of residues with the highest predicted flexibility probability were selected), the model applies column-wise masking and deep sampling to the MSA, generating diverse sub-MSAs that encode distinct co-evolutionary information. These sub-MSAs are then fed into AlphaFold2^2^ to predict multiple conformational states, which provide diverse distance maps to guide the optimization in M-DeepAssembly2. The detailed implementation of the network can be found in our previously developed AlphaFlex^20^ method.

### Multi-objective protein conformation sampling algorithm

The distance maps obtained from the flexible residue prediction network^20^ and the DeepAssembly^26^ method were employed as energy functions in a multi-objective optimization model. The Pareto solutions derived from these distinct energy functions can be described by the following equation^36^.

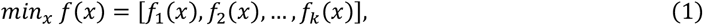

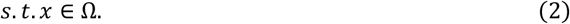

where *f*_*i*_(*x*), *iϵ*{1,2, …, *k*} is the *i*-th energy function, *k* represents the total number of energy functions, and *x* is the decision vector in the decision space Ω. For two conformation *x*^*a*^ and *x*^*b*^ in the conformational space, for all energy function *f*_*i*_(*x*), if *f*_*j*_(*x*^*a*^) ≤ *f*_*j*_(*x*^*b*^), ∀*j* ∈ {1,2, …, *m*} and *f*_*q*_(*x*^*a*^) < *f*_*q*_(*x*^*b*^), ∃ *q* ∈ {1,2, …, *m*}, then *x*^*a*^ is said to dominate *x*^*b*^(donated as *x*^*a*^ ≺ *x*^*b*^). The set of all non-dominated solutions constitutes the Pareto solution set^19^.

### CRediT authorship contribution statements

**Xinyue Cui:** Writing-review & editing, Writing-original draft, Visualization, Software, Methodology, Data curation. **Lingyu Ge:** Writing-review & editing, Software, Methodology. **Xinguang Yang**: Writing-review & editing, Software, Methodology. **Xuhui Li**: Writing-review & editing, Software, Methodology. **Guijun Zhang**: Writing-review & editing, Validation, Supervision, Resources, Project administration, Methodology, Funding acquisition, Conceptualization.

## Acknowledgements

This work was supported by the National Key R&D Program of China [2022ZD0115103], the National Nature Science Foundation of China [62173304], the “Pioneer” and “Leading Goose” R&D Program of Zhejiang [2025C01190], and the Zhejiang Province High-level Talent Special Support Program [2023R5248].

## Appendix A. Supplementary material

Supplementary material to this article can be found at http://zhanglab-bioinf.com/M-DeepAssembly2/.

## Reference

1 Roy, A., Hua, D. P., Post, C. B. (2016). Analysis of multidomain protein dynamics. Journal of Chemical Theory and Computation. 12, 274–280. 10.1021/acs.jctc.5b00796.

2 Jumper, J., Evans, R., Pritzel, A., Green, T., Figurnov, M., Ronneberger, O., Tunyasuvunakool, K., Bates, R., Žídek, A., Potapenko, A. (2021). Highly accurate protein structure prediction with AlphaFold. Nature. 596, 583–589. 10.1038/s41586-021-03819-2.

3 Abramson, J., Adler, J., Dunger, J., Evans, R., Green, T., Pritzel, A., Ronneberger, O., Willmore, L., Ballard, A. J., Bambrick, J. (2024). Accurate structure prediction of biomolecular interactions with AlphaFold 3. Nature. 630, 493–500. 10.1038/s41586-024-07487-w.

4 Baek, M., DiMaio, F., Anishchenko, I., Dauparas, J., Ovchinnikov, S., Lee, G. R., Wang, J., Cong, Q., Kinch, L. N., Schaeffer, R. D. (2021). Accurate prediction of protein structures and interactions using a three-track neural network. Science. 373, 871–876. 10.1126/science.abj8754.

5 Lin, Z., Akin, H., Rao, R., Hie, B., Zhu, Z., Lu, W., Smetanin, N., Verkuil, R., Kabeli, O., Shmueli, Y. (2023). Evolutionary- scale prediction of atomic-level protein structure with a language model. Science. 379, 1123–1130. 10.1126/science.ade2574.

6 Xia, Y., Pu, Y., Wang, S., Zhuang, J., Liu, D., Hou, M., Zhang, G. (2025). DeepAssembly2: A Web Server for Protein Complex Structure Assembly Based on Domain-Domain Interactions. Journal of Molecular Biology. 169128. 10.1016/j.jmb.2025.169128.

7 Wüthrich, K. (1986). NMR with proteins and nucleic acids. Europhysics News. 17, 11–13. 10.1051/epn/19861701011.

8 Cheng, Y. (2015). Single-particle cryo-EM at crystallographic resolution. Cell. 161, 450–457. 10.1016/j.cell.2015.03.049.

9 Smyth, M.,Martin, J. (2000). x Ray crystallography. Molecular Pathology. 53, 8. 10.1136/mp.53.1.8.

10 Gaalswyk, K., Muniyat, M. I., MacCallum, J. L. (2018). The emerging role of physical modeling in the future of structure determination. Current Opinion in Structural Biology. 49, 145–153. 10.1016/j.sbi.2018.03.005.

11 Cui, X., Ge, L., Chen, X., Lv, Z., Wang, S., Zhou, X., Zhang, G. (2025). Beyond static structures: protein dynamic conformations modeling in the post-AlphaFold era. Briefings in Bioinformatics. 26, bbaf340. 10.1093/bib/bbaf340.

12 Wayment-Steele, H. K., Ojoawo, A., Otten, R., Apitz, J. M., Pitsawong, W., Hömberger, M., Ovchinnikov, S., Colwell, L., Kern, D. (2024). Predicting multiple conformations via sequence clustering and AlphaFold2. Nature. 625, 832–839. 10.1038/s41586-023-06832-9.

13 Kalakoti, Y.,Wallner, B. (2025). AFsample2 predicts multiple conformations and ensembles with AlphaFold2. Communications Biology. 8, 373. 10.1038/s42003-025-07791-9.

14 Jing, B., Berger, B., Jaakkola, T. (2024). AlphaFold meets flow matching for generating protein ensembles. Arxiv preprint 2402.04845. 10.48550/arXiv.2402.04845.

15 Zheng, S., He, J., Liu, C., Shi, Y., Lu, Z., Feng, W., Ju, F., Wang, J., Zhu, J., Min, Y. (2024). Predicting equilibrium distributions for molecular systems with deep learning. Nature Machine Intelligence. 6, 558–567. 10.1038/s42256-024-00837-3.

16 Lewis, S., Hempel, T., Jiménez-Luna, J., Gastegger, M., Xie, Y., Foong, A. Y., Satorras, V. G., Abdin, O., Veeling, B. S., Zaporozhets, I. (2025). Scalable emulation of protein equilibrium ensembles with generative deep learning. Science. eadv9817. 10.1126/science.adv9817.

17 Peng, C., Zhou, X., Liu, J., Hou, M., Li, S. Z., Zhang, G. (2024). Multiple conformational states assembly of multidomain proteins using evolutionary algorithm based on structural analogues and sequential homologues. Fundamental Research. 10.1016/j.fmre.2024.05.003.

18 Hou, M., Jin, S., Cui, X., Peng, C., Zhao, K., Song, L., Zhang, G. (2024). Protein multiple conformation prediction using multi-objective evolution algorithm. Interdisciplinary Sciences: Computational Life Sciences. 16, 519–531. 10.1007/s12539-023-00597-5.

19 Cui, X., Xia, Y., Hou, M., Zhao, X., Wang, S., Zhang, G. (2025). M-DeepAssembly: enhanced DeepAssembly based on multi-objective multi-domain protein conformation sampling. BMC Bioinformatics. 26, 120. 10.1186/s12859-025-06131-2.

20 Ge, L., Cui, X., Zhao, K., Zhou, X., Zhang, Y., Zhang, G. (2025). AlphaFlex: Accuracy modeling of protein multiple conformations via predicted flexible residues. Biorxiv. 2025.2007. 2011.664327. 10.1101/2025.07.11.664327.

21 Yu, Z., Peng, C., Liu, J., Zhang, B., Zhou, X., Zhang, G. (2022). DomBpred: protein domain boundary prediction based on domain-residue clustering using inter-residue distance. IEEE/ACM Transactions on Computational Biology and Bioinformatics. 20, 912–922. 10.1109/TCBB.2022.3175905.

22 Rao, R. M., Liu, J., Verkuil, R., Meier, J., Rives, A. (2021). MSA transformer. ICML. 10.1101/2021.02.12.430858.

23 Henikoff, S.,Henikoff, J. G. (1992). Amino acid substitution matrices from protein blocks. Proceedings of the National Academy of Sciences. 89, 10915–10919. 10.1073/pnas.89.22.10915.

24 Yang, J., Anishchenko, I., Park, H., Peng, Z., Ovchinnikov, S., Baker, D. (2020). Improved protein structure prediction using predicted interresidue orientations. Proceedings of the National Academy of Sciences. 117, 1496–1503. 10.1073/pnas.1914677117.

25 Guo, S., Liu, J., Zhou, X., Zhang, G. (2022). DeepUMQA: ultrafast shape recognition-based protein model quality assessment using deep learning. Bioinformatics. 38, 1895–1903. 10.1093/bioinformatics/btac056.

26 Xia, Y., Zhao, K., Liu, D., Zhou, X., Zhang, G. (2023). Multi-domain and complex protein structure prediction using inter- domain interactions from deep learning. Communications Biology. 6, 1221. 10.1038/s42003-023-05610-7.

27 Monzon, A. M., Rohr, C. O., Fornasari, M. S., Parisi, G. (2016). CoDNaS 2.0: a comprehensive database of protein conformational diversity in the native state. Database. 2016, baw038. 10.1093/database/baw038.

28 Zhang, Y.,Skolnick, J. (2004). Scoring function for automated assessment of protein structure template quality. Proteins: Structure, Function, and Bioinformatics. 57, 702–710. 10.1002/prot.20264.

29 Liu, D., Zhao, X., Zhang, T., Xie, L., Ye, E., Liang, F., Wang, H., Zhang, G. (2025). MViewEMA: Efficient Global Accuracy Estimation for Protein Complex Structural Models Using Multi-View Representation Learning. bioRxiv. 2025.2007. 2025.666906. 10.1101/2025.07.25.666906.

30 Liu, D., Liu, J., Wang, H., Liang, F., Zhang, G. (2025). DeepUMQA-X: Comprehensive and insightful estimation of model accuracy for protein single-chain and complex. Nucleic Acids Research. gkaf380. 10.1093/nar/gkaf380.

31 Shumilin, I. A., Bauerle, R., Wu, J., Woodard, R. W., Kretsinger, R. H. (2004). Crystal structure of the reaction complex of 3-deoxy-D-arabino-heptulosonate-7-phosphate synthase from Thermotoga maritima refines the catalytic mechanism and indicates a new mechanism of allosteric regulation. Journal of Molecular Biology. 341, 455–466. 10.1016/j.jmb.2004.05.077.

32 Piazza, M., Taiakina, V., Dieckmann, T., Guillemette, J. G. (2017). Structural consequences of calmodulin EF hand mutations. Biochemistry. 56, 944–956. 10.1021/acs.biochem.6b01296.

33 Suzek, B. E., Wang, Y., Huang, H., McGarvey, P. B., Wu, C. H., Consortium, U. (2015). UniRef clusters: a comprehensive and scalable alternative for improving sequence similarity searches. Bioinformatics. 31, 926–932. 10.1093/bioinformatics/btu739.

34 Mirdita, M., Von Den Driesch, L., Galiez, C., Martin, M. J., Söding, J., Steinegger, M. (2017). Uniclust databases of clustered and deeply annotated protein sequences and alignments. Nucleic Acids Research. 45, D170–D176. 10.1093/nar/gkw1081.

35 Steinegger, M.,Söding, J. (2018). Clustering huge protein sequence sets in linear time. Nature Communications. 9, 2542. 10.1038/s41467-018-04964-5.

36 Zitzler, E.,Thiele, L. (2002). Multiobjective evolutionary algorithms: a comparative case study and the strength Pareto approach. IEEE transactions on Evolutionary Computation. 3, 257–271. 10.1109/4235.797969.

