## Supplemental Tables and Figures for "M-DeepAssembly2: A Web Server for Predicting Multiple Conformations of Multi-domain Proteins Using Deep Learning"

**This file includes:**

- Supplementary Tables S1 to S5
- Supplementary Figures S1 to S4

### Supplementary Tables

**Supplementary Table S1.** Performance of M-DeepAssembly2 on 14 Apo/Holo protein pairs (Apo state).

| Targets | M-DeepAssembly2 | | AF-Cluster | | AlphaFLOW | |
|---|---|---|---|---|---|---|
| | TM-score | RMSD(Å) | TM-score | RMSD(Å) | TM-score | RMSD(Å) |
| 1cfcA | 0.767 | 2.792Å | 0.627 | 4.191Å | 0.737 | 2.987Å |
| 1dmoA | 0.887 | 1.814Å | 0.603 | 5.299Å | 0.796 | 2.564Å |
| 1evkA | 1.000 | 0.075Å | 0.983 | 0.968Å | 0.973 | 1.225Å |
| 1g6wD | 1.000 | 0.045Å | 0.939 | 1.803Å | 0.954 | 1.389Å |
| 1hw1B | 0.995 | 0.407Å | 0.746 | 4.264Å | 0.923 | 1.689Å |
| 1jejA | 0.998 | 0.322Å | 0.636 | 8.934Å | 0.969 | 1.278Å |
| 1jfjA | 0.614 | 6.168Å | 0.422 | 10.820Å | 0.455 | 7.019Å |
| 1k5hA | 0.991 | 0.677Å | 0.901 | 4.776Å | 0.923 | 4.475Å |
| 1njgB | 0.994 | 0.458Å | 0.950 | 1.411Å | 0.954 | 1.326Å |
| 1tjdA | 0.893 | 2.067Å | 0.922 | 1.758Å | 0.929 | 1.724Å |
| 1vr6A | 0.999 | 0.120Å | 0.951 | 1.694Å | 0.926 | 2.045Å |
| 1w0jE | 0.994 | 0.582Å | 0.967 | 1.507Å | 0.976 | 1.241Å |
| 2cg7A | 0.735 | 2.746Å | 0.838 | 1.745Å | 0.923 | 1.061Å |
| 2rcsH | 0.878 | 2.126Å | 0.699 | 4.348Å | 0.674 | 4.472Å |
| **Average** | **0.910** | **1.457Å** | **0.799** | **3.892**Å | **0.865** | **2.464**Å |

**Supplementary Table S2.** Performance of M-DeepAssembly2 on 14 Apo/Holo protein pairs (Holo state).

| Targets | M-DeepAssembly2 | | AF-Cluster | | AlphaFLOW | |
|---|---|---|---|---|---|---|
| | TM-score | RMSD(Å) | TM-score | RMSD(Å) | TM-score | RMSD(Å) |
| 5dowA | 0.595 | 5.841Å | 0.719 | 3.385Å | 0.658 | 4.078Å |
| 3clnA | 0.932 | 1.276Å | 0.813 | 2.340Å | 0.650 | 4.495Å |
| 1evlA | 0.982 | 1.049Å | 0.982 | 1.023Å | 0.967 | 1.378Å |
| 1k0bC | 0.986 | 0.723Å | 0.958 | 1.446Å | 0.955 | 1.368Å |
| 1h9gA | 0.957 | 1.316Å | 0.734 | 4.513Å | 0.934 | 1.614Å |
| 1jG6A | 0.992 | 0.692Å | 0.645 | 9.016Å | 0.960 | 1.492Å |
| 1jfkA | 0.675 | 3.483Å | 0.520 | 6.809Å | 0.502 | 8.870Å |
| 1q0qA | 0.978 | 1.146Å | 0.977 | 1.138Å | 0.962 | 1.471Å |
| 1njfA | 0.978 | 1.021Å | 0.983 | 0.776Å | 0.956 | 1.407Å |
| 1eejB | 0.841 | 2.683Å | 0.937 | 1.508Å | 0.921 | 1.900Å |
| 1rzmA | 0.919 | 2.171Å | 0.902 | 2.843Å | 0.964 | 1.380Å |
| 1e1rF | 0.910 | 2.516Å | 0.911 | 2.613Å | 0.937 | 2.060Å |
| 2cg6A | 0.646 | 4.408Å | 0.570 | 3.702Å | 0.568 | 5.492Å |
| 1aj7H | 0.694 | 4.189Å | 0.747 | 3.677Å | 0.631 | 5.078Å |
| **Average** | **0.863** | **2.322Å** | **0.806** | **3.277Å** | **0.826** | **3.006Å** |

**Supplementary Table S3.** Structural differences between 14 Apo/Holo pairs.

| Apo | Holo | TM-score | RMSD(Å) |
|---|---|---|---|
| 1cfcA | 5dowA | 0.377 | 9.247Å |
| 1dmoA | 3clnA | 0.396 | 13.274Å |
| 1evkA | 1evlA | 0.958 | 1.635Å |
| 1g6wD | 1k0bC | 0.962 | 1.360Å |
| 1hw1B | 1h9gA | 0.930 | 1.672Å |
| 1jejA | 1jG6A | 0.925 | 2.099Å |
| 1jfjA | 1jfkA | 0.466 | 14.727Å |
| 1k5hA | 1q0qA | 0.859 | 4.926Å |
| 1njgB | 1njfA | 0.954 | 1.332Å |
| 1tjdA | 1eejB | 0.887 | 2.078Å |
| 1vr6A | 1rzmA | 0.815 | 10.107Å |
| 1w0jE | 1e1rF | 0.821 | 3.888Å |
| 2cg7A | 2cg6A | 0.553 | 6.233Å |
| 2rcsH | 1aj7H | 0.588 | 7.207Å |
| **Average** | | **0.749** | **5.699Å** |

**Supplementary Table S4.** Performance of M-DeepAssembly2 on 14 static multi-domain proteins.

| Targets | M-DeepAssembly2 | | M-DeepAssembly | | AlphaFold2 | | AlphaFold3 | |
|---|---|---|---|---|---|---|---|---|
| | **TM-score** | **RMSD(Å)** | **TM-score** | **RMSD(Å)** | **TM-score** | **RMSD(Å)** | **TM-score** | **RMSD(Å)** |
| 1oqyA | 0.224 | 40.172Å | 0.226 | 28.589Å | 0.226 | 45.946Å | 0.223 | 50.118Å |
| 1zzaA | 0.415 | 18.245Å | 0.392 | 15.568Å | 0.319 | 18.923Å | 0.316 | 22.563Å |
| 2jt3A | 0.402 | 13.297Å | 0.412 | 12.363Å | 0.440 | 11.361Å | 0.469 | 9.376Å |
| 2kn6A | 0.457 | 17.980Å | 0.459 | 15.481Å | 0.459 | 15.940Å | 0.474 | 13.615Å |
| 2kr0A | 0.311 | 32.345Å | 0.306 | 35.816Å | 0.304 | 34.564Å | 0.298 | 32.754Å |
| 2l6lA | 0.396 | 13.059Å | 0.387 | 12.415Å | 0.388 | 11.816Å | 0.394 | 14.029Å |
| 2looA | 0.343 | 7.556Å | 0.399 | 6.453Å | 0.324 | 11.645Å | 0.314 | 13.816Å |
| 2lqnA | 0.542 | 21.004Å | 0.371 | 21.187Å | 0.362 | 24.702Å | 0.350 | 21.668Å |
| 4d1eA | 0.378 | 99.304Å | 0.356 | 100.270Å | 0.426 | 99.680Å | 0.437 | 100.132Å |
| 5t0gC | 0.583 | 14.674Å | 0.432 | 15.265Å | 0.430 | 7.583Å | 0.613 | 10.732Å |
| 5tp5A | 0.329 | 11.535Å | 0.380 | 11.494Å | 0.372 | 11.040Å | 0.352 | 13.094Å |
| 5tp6A | 0.662 | 3.894Å | 0.335 | 11.439Å | 0.380 | 12.917Å | 0.421 | 16.528Å |
| 6kvgA | 0.681 | 16.207Å | 0.453 | 10.085Å | 0.443 | 18.472Å | 0.495 | 18.739Å |
| 7dmeA | 0.451 | 26.646Å | 0.442 | 25.124Å | 0.446 | 23.906Å | 0.457 | 22.278Å |
| **Average** | **0.441** | **23.994Å** | **0.382** | **22.968Å** | **0.380** | **24.893Å** | **0.401** | **25.674Å** |

**Supplementary Table S5.** Performance of M-DeepAssembly2* and M-DeepAssembly2 on 14 multi-domain proteins.

| **Targets** | **M-DeepAssembly2*** | | **M-DeepAssembly2** | |
|---|---|---|---|---|
| | **TM-score** | **RMSD(Å)** | **TM-score** | **RMSD(Å)** |
| 1oqyA | 0.261 | 22.328Å | 0.224 | 40.172Å |
| 1zzaA | 0.447 | 14.136Å | 0.415 | 18.245Å |
| 2jt3A | 0.602 | 5.458Å | 0.402 | 13.297Å |
| 2kn6A | 0.528 | 9.770Å | 0.457 | 17.980Å |
| 2kr0A | 0.390 | 30.300Å | 0.311 | 32.345Å |
| 2l6lA | 0.455 | 8.632Å | 0.396 | 13.059Å |
| 2looA | 0.440 | 6.685Å | 0.343 | 7.556Å |
| 2lqnA | 0.550 | 14.630Å | 0.542 | 21.004Å |
| 4d1eA | 0.489 | 93.223Å | 0.378 | 99.304Å |
| 5t0gC | 0.665 | 6.491Å | 0.583 | 14.674Å |
| 5tp5A | 0.531 | 6.215Å | 0.329 | 11.535Å |
| 5tp6A | 0.673 | 3.822Å | 0.662 | 3.894Å |
| 6kvgA | 0.685 | 15.918Å | 0.681 | 16.207Å |
| 7dmeA | 0.466 | 12.443Å | 0.451 | 26.646Å |
| **Average** | **0.513** | **17.861Å** | **0.441** | **23.994Å** |

*Note: M-DeepAssembly2* refers to the predicted structure exhibiting the highest structural similarity to the native structure. M-DeepAssembly3 denotes the best model selected using our in-house developed MViewEMA model quality assessment algorithm.*

### Supplementary Figures

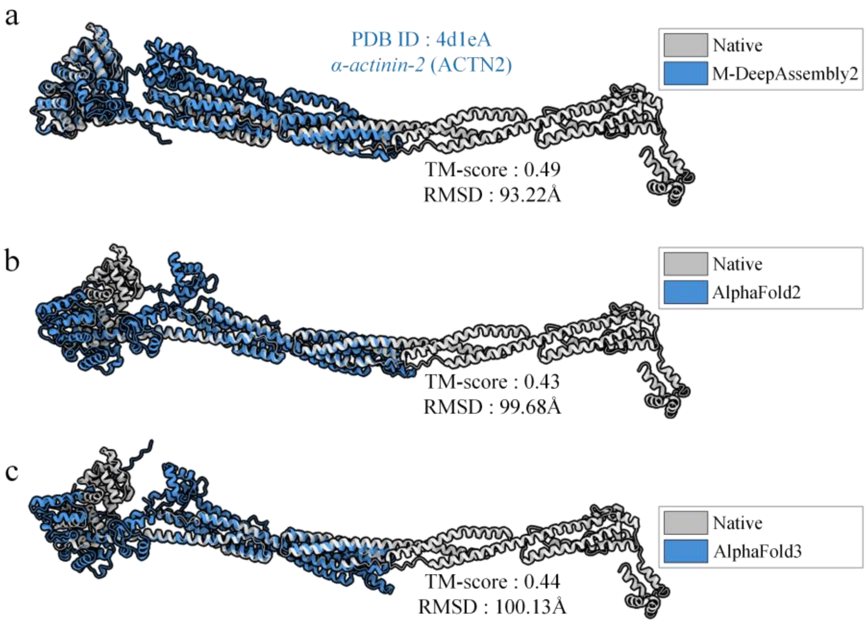


**Supplementary Figure S1.** Prediction results of protein 4d1eA using different methods. **a,** Comparison of M-DeepAssembly2 predictions with the native structure. **b,** Comparison of AlphaFold2 predictions with the native structure. **c,** Comparison of AlphaFold3 predictions with the native structure.

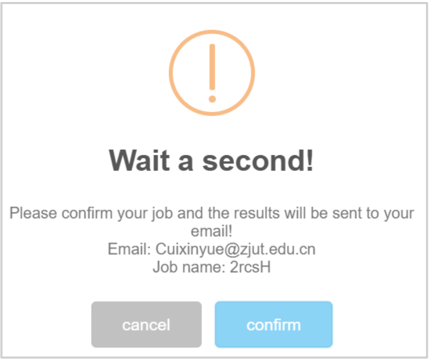


**Supplementary Figure S2.** A confirmation dialog for task submission, which displays the user's email and the task name.

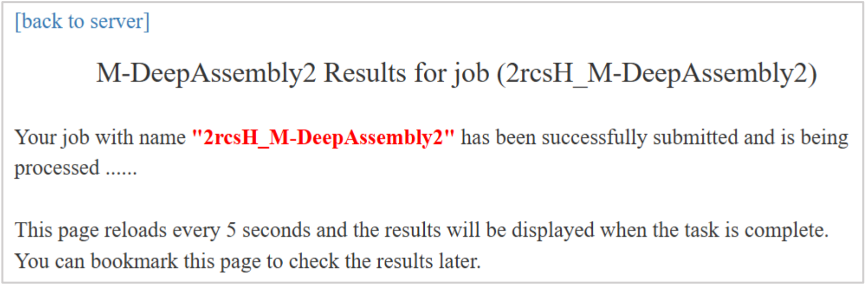


**Supplementary Figure S3.** Diagram of the page redirection after successful task submission.

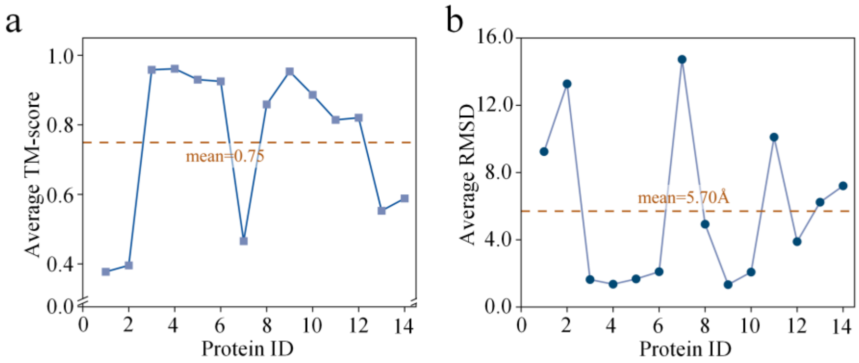


**Supplementary Figure S4.** Conformational similarity analysis of 14 Apo/Holo protein pairs. **a**, TM-scores among the 14 Apo/Holo pairs with an average value of 0.75. **b**, RMSD values among the 14 Apo/Holo pairs with an average value of 5.70 Å.
